# Semi-Automatic Detection of Errors in Genome-Scale Metabolic Models

**DOI:** 10.1101/2024.06.24.600481

**Authors:** Devlin C. Moyer, Justin Reimertz, Daniel Segrè, Juan I. Fuxman Bass

## Abstract

**Background:** Genome-Scale Metabolic Models (GSMMs) are used for numerous tasks requiring computational estimates of metabolic fluxes, from predicting novel drug targets to engineering microbes to produce valuable compounds. A key limiting step in most applications of GSMMs is ensuring their representation of the target organism’s metabolism is complete and accurate. Identifying and visualizing errors in GSMMs is complicated by the fact that they contain thousands of densely interconnected reactions. Furthermore, many errors in GSMMs only become apparent when considering pathways of connected reactions collectively, as opposed to examining reactions individually.

**Results:** We present Metabolic Accuracy Check and Analysis Workflow (MACAW), a collection of algorithms for detecting errors in GSMMs. The relative frequencies of errors we detect in manually curated GSMMs appear to reflect the different approaches used to curate them. Changing the method used to automatically create a GSMM from a particular organism’s genome can have a larger impact on the kinds of errors in the resulting GSMM than using the same method with a different organism’s genome. Our algorithms are particularly capable of identifying errors that are only apparent at the pathway level, including loops, and nontrivial cases of dead ends.

**Conclusions:** MACAW is capable of identifying inaccuracies of varying severity in a wide range of GSMMs. Correcting these errors can measurably improve the predictive capacity of a GSMM. The relative prevalence of each type of error we identify in a large collection of GSMMs could help shape future efforts for further automation of error correction and GSMM creation.

## Background

Genome-Scale Metabolic Models (GSMMs) are formal representations of metabolism that enable mathematical predictions of fluxes through a cell’s metabolic network. GSMMs have been widely used to simulate the metabolic phenotypes of cells and metabolic interactions between groups of cells [1–4]. GSMMs have been used in contexts as varied as predicting novel drug targets for various human diseases [5–13], characterizing the metabolic differences between different human cell (sub)types [14–20], and helping engineer microbes to produce commercially and/or medically valuable compounds [3,21–25]. GSMMs represent metabolic networks as stoichiometric matrices, where each row in the matrix corresponds to a single metabolite, each column represents a reaction, and each entry contains the stoichiometric coefficient of that row’s metabolite in that column’s reaction. These stoichiometric matrices are also typically annotated with information about which genes encode the enzymes (or subunits of enzyme complexes) that catalyze each reaction in the network, which enable GSMMs to serve as platforms for integrating many kinds of data, including transcriptomics, proteomics, metabolomics, thermodynamic, and kinetic data [12,20,26–28]. In particular, GSMMs are most commonly used to predict steady-state metabolic fluxes through all of the reactions they contain, and integrating ‘omics data can help tailor these predictions to specific biological contexts [4,19,29–34].

While GSMMs have been developed for many different organisms and cell types, the accuracy and completeness of GSMMs in general remains highly variable, despite many efforts to systematically test for and correct errors. Even GSMMs that were extensively manually curated and formally published can contain numerous and varied errors, which limit their practical applicability [35–38]. These errors can include reactions with inaccurate stoichiometric coefficients or reversibilities, inaccurate associations between reactions and genes, duplicate reactions, reactions that are incapable of sustaining steady-state fluxes, and reactions capable of infinite or otherwise implausible fluxes [35–38]. Some of these are human errors, introduced during manual construction or curation of GSMMs, while others arise from flaws in the heuristics used by automated construction or curation tools [39]. Additionally, new experiments occasionally demonstrate that, for instance, a particular organism or enzyme is incapable of catalyzing a particular reaction that it was initially thought to catalyze, rendering a GSMM that appeared to be accurate when it was first constructed appear less accurate over time. Many existing tools for automatically detecting and correcting errors in GSMMs, such as Meneco and fastGapFill [40,41], primarily focus on reactions that are incapable of sustaining steady-state fluxes due to the presence of dead-end metabolites that can only be produced and never consumed or vice versa [42–47]. While these so-called “gap-filling” algorithms play a useful role in the creation of new GSMMs, especially for poorly studied organisms, many are prone to introducing new errors as they attempt to connect the dead-ends to the rest of the network [47–49]. Other tools focus on identifying and correcting loops of reactions that are capable of sustaining arbitrarily large (and thus thermodynamically infeasible) cycles of flux [50]. Unfortunately, these tools are also prone to blocking fluxes through reactions representing electron transport chains and ATP synthases, which predictably renders the GSMMs they modify noticeably less capable of modeling realistic metabolic phenotypes [50].

A few existing tools for curating GSMMs, such as MEMOTE [51] and ErrorTracer [52], take a different approach: rather than automatically fixing the errors they identify, they focus on highlighting potentially problematic reactions, leaving it to the user to manually investigate each reaction and decide what the appropriate fix is. Even though they do not remove the need for manual curation, prioritizing where to look for problems becomes a non-trivial task when curating GSMMs that have thousands of reactions. In this vein, we introduce Metabolic Accuracy Check and Analysis Workflow (MACAW), a collection of new algorithms for detecting errors, especially pathway-level errors, in GSMMs (Fig. 1A). In addition to highlighting reactions, MACAW also connects highlighted reactions into networks to help users visualize pathway-level errors. We demonstrate MACAW’s utility by showcasing several errors that it highlights in a selection of manually-curated GSMMs [53–55], as well as broader trends it highlights in large collections of automatically generated GSMMs [56,57]. In particular, we show that we were able to identify and correct errors with 615 of the 13,083 reactions in version 1.15 of Human-GEM by following up on reactions highlighted by MACAW, and that correcting these errors improves Human-GEM’s ability to predict the impact of several knockouts in the lipoic acid biosynthesis pathway.

**Figure 1.**
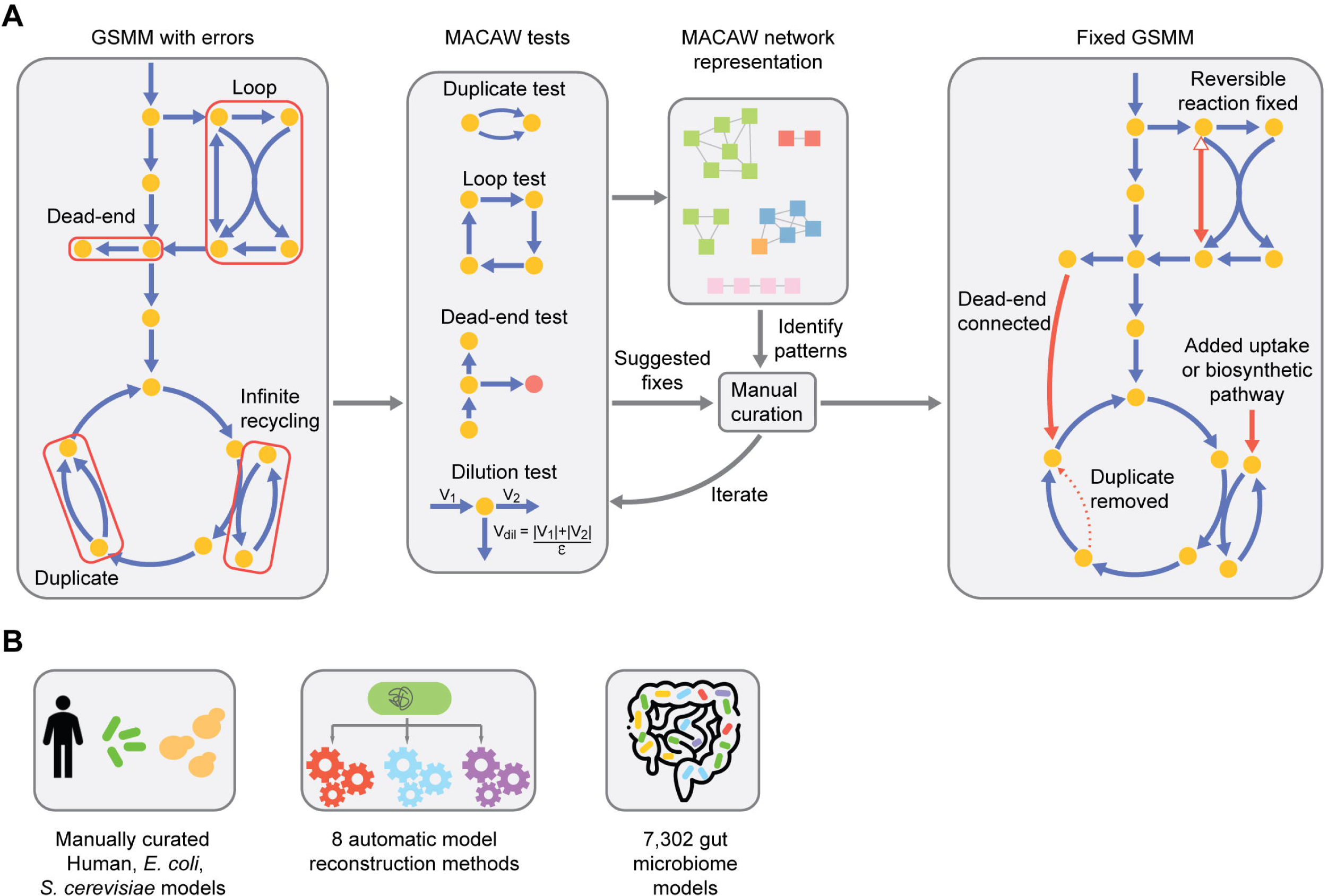
Overview of MACAW. (**A**) Schematic showing the application of MACAW to a toy GSMM containing errors. The duplicate, loop, dead-end, and dilution tests are applied to the GSMM. MACAW then suggests potential fixes and provides a network visualization connecting flagged reactions. This information can be used iteratively to identify patterns and manually curate the GSMM. Fixed reactions are depicted by red arrows. (**B**) GSMMs to which MACAW was applied in this manuscript.

## Results

### Metabolic Accuracy Check and Analysis Workflow (MACAW)

MACAW contains four independent but complementary tests that can be run on an arbitrary GSMM to highlight reactions that are potentially inaccurate: the dead-end test, dilution test, duplicate test, and loop test (Fig. 1A). Some tests are extensions of tests that have already been published elsewhere [51,52], while others involve novel algorithms.

The most innovative test is the dilution test, which primarily identifies metabolites that can only be recycled into each other and never produced from external sources or secreted (see “Methods”). While many metabolites function as cofactors, repeatedly being interconverted between two or more forms (e.g. ATP and ADP), all cells must have biosynthetic pathways to create these cofactors or uptake pathways in order to counter dilution of cofactors due to cellular growth and division and/or loss to side reactions (e.g., with reactive oxygen species) [58,59]. Since the most common application of GSMMs is to use them to predict steady-state fluxes through reactions, it can be easy to miss that a particular GSMM is incapable of *net* production (as opposed to recycling) of a particular cofactor. This can be especially problematic if one is trying to use a GSMM to study disorders of cofactor metabolism, for instance. The dilution test highlights these kinds of errors by imposing so-called “dilution” constraints on the GSMM and identifying which reactions are able to sustain non-zero fluxes before but not after imposing dilution constraints [60,61]. Imposing dilution constraints on a GSMM involves creating a new reaction for each metabolite that consumes it and produces no products (representing its “dilution” away from the cell being modeled and/or its loss to side-reactions) and then constraining the flux through each dilution reaction to be some fraction of the sum of the fluxes through all other reactions the corresponding metabolite participates in (see “Methods”). To our knowledge, dilution constraints have never been used to systematically identify errors in GSMMs, and we show below that we are able to identify and correct several missing reactions and other errors in cofactor metabolism in Human-GEM, which improves Human-GEM’s ability to accurately predict the impact of knockouts of several genes.

A common way to use GSMMs is to predict steady-state fluxes through all the reactions they contain, and a common problem encountered when doing so is loops of reactions that can sustain arbitrarily large thermodynamically infeasible cyclic fluxes [38,50,62,63]. While the existence of such loops is not always attributable to mistakes in the construction of the underlying network, it is occasionally the case that correcting errors in a GSMM can reduce the number of loops. For example, a pair of duplicate reactions oriented in opposite directions that could be replaced by a single reversible reaction, or a reversible reaction that is known to primarily carry fluxes in a single direction in the organism being modeled. The loop test identifies all reactions that are capable of non-zero fluxes when all exchange (uptake/import and/or secretion/export) reactions in the GSMM are blocked. While a number of existing tools do the same, the loop test in MACAW also groups all reactions it identifies into distinct loops (see “Methods”). This significantly streamlines the process of investigating whether or not any inaccuracies in the GSMM are responsible for each loop, given that many GSMMs contain hundreds of reactions involved in such loops.

GSMMs often contain groups of identical or near-identical reactions that correspond to a single real-life reaction, due to errors in their construction and/or curation. These duplicate reactions can sometimes create infinite loops of flux between the two duplicate reactions and/or make it difficult to constrain fluxes through reactions based on expression levels of the enzymes that catalyze them. The duplicate test highlights all groups of reactions that involve the same set of metabolites, but may have different stoichiometric coefficients, reversibilities, and/or associated genes. Existing GSMM curation tools, such as MEMOTE, also have similar duplicate tests, but the MACAW duplicate test flags a broader range of duplicates. In particular, the MEMOTE [51] duplicate test only tests reactions that have an International Chemical Identifier (InChI) [64] associated with all metabolites, while the duplicate test in MACAW has no such requirement and is thus capable of identifying potential errors that existing tools would miss.

Many GSMMs contain many reactions that form dead-ends, e.g. pathways that end with a metabolite that can only be produced but never consumed, and are thus incapable of sustaining steady-state fluxes. MACAW’s dead-end test highlights reactions that produce metabolites that cannot be consumed by any other reaction in the GSMM or vice versa. While several existing tools include superficially similar tests, they are implemented in significantly different ways than the dead-end test in MACAW. Many existing implementations report all metabolites that are produced or are consumed by all reactions they participate in without reporting which reactions are incapable of steady-state fluxes due to these dead-ends. Others use Flux Variability Analysis (FVA) [65] to identify all reactions that are incapable of non-zero steady-state fluxes without noting which dead-end metabolites are responsible for blocking each reaction. In both cases, existing dead-end tests rarely provide any information on how the metabolites or reactions that they highlight connect to each other. The primary innovation of the dead-end test in MACAW is that it connects each dead-end metabolite it identifies to all reactions that it prevents from carrying flux, tracing out distinct dead-end pathways within the tested GSMM. These pathways facilitate correcting each dead-end by providing relevant context and clearly showing when a group of multiple dead-end reactions can all be fixed by correcting a single dead-end (e.g., by adding a new reaction that consumes a metabolite that was previously only capable of being produced). As discussed in more detail below, our approach also identifies certain dead-end pathways that many previous approaches would not have found.

### Application of MACAW to manually curated human, yeast, and *E. coli* GSMMs

We first ran MACAW on three of the most extensively curated and broadly used GSMMs available: Human-GEM version 1.15 [53], yeast-GEM version 9.0.0 [54] (for *Saccharomyces cerevisiae*), and iML1515 [55] (one of the most recently published GSMMs for *Escherichia coli*) (Fig. 1B). The proportion of reactions flagged by any of the MACAW tests varied between the three models: 4,783 of 13,073 (37%) of reactions were flagged by any test in Human-GEM, 2,061 of 4,130 (50%) in yeast-GEM, and 1,406 of 2,712 (52%) in iML1515 (Fig. 2). It is important to note that the tests in MACAW are designed to flag reactions that are worth examining for potential errors as well as reactions that may simply be close network neighbor of erroneous reactions, so the percentages listed here should not be interpreted as the percentage of reactions in each GSMM that contain one or more errors. Nonetheless, these results suggest that many errors remain to be corrected even in these highly curated GSMMs.

**Figure 2.**
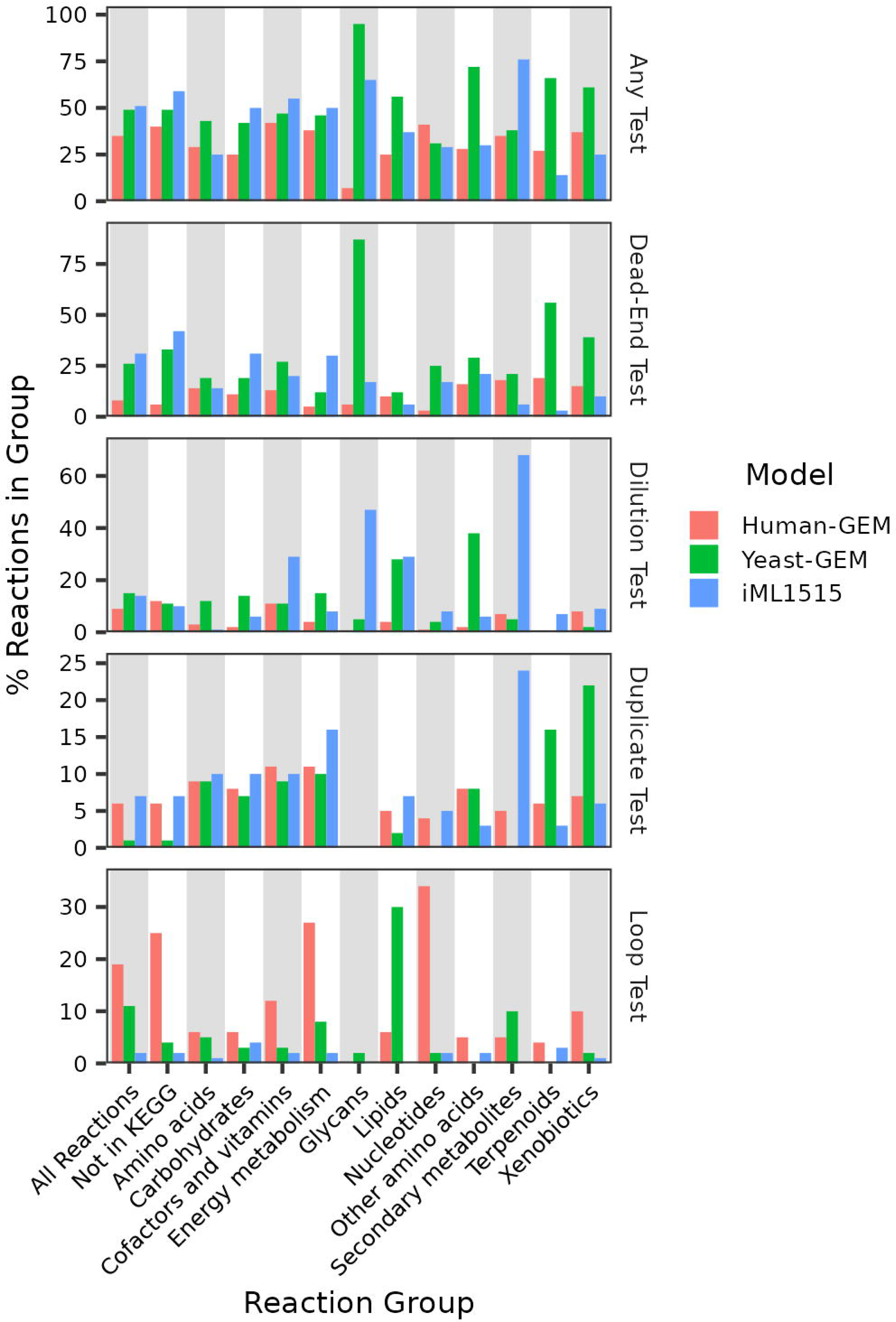
Proportions of reactions flagged by each test in manually curated GSMMs. Reactions grouped according to the second-highest levels of the KEGG functional ortholog hierarchy containing the genes associated with each reaction; see “Methods”. Individual reactions may be associated with more than one group of KEGG functional orthologs. Names of KEGG functional ortholog groups have been abbreviated.

To investigate whether the different tests flag reactions in different parts of each metabolic network, we categorized each reaction using the KEGG Orthology database [66] (Fig. 2). Overall, the tests flag a diverse set of reactions in all three GSMMs, including extensively studied areas of metabolism, such as amino acid, carbohydrate, and energy metabolism. The dead-end test and the duplicate test flag similar proportions of reactions in most classes of reactions across the three GSMMs, suggesting that the errors they target are not associated with any particular kind of reactions or attributable to differences in how the GSMMs were curated. Conversely, the dilution and loop tests flag variable proportions of reactions both between reaction categories and GSMMs, suggesting that the errors they target might be associated with particular kinds of reactions and/or differences in the ways that the three GSMMs were curated.

In particular, the loop test flagged a significantly larger proportion of energy and nucleotide metabolism reactions in Human-GEM than in yeast-GEM or iML1515, as well as a larger proportion of lipid metabolism in yeast-GEM than Human-GEM or iML1515 (Fig. 2). This may be attributable to differences in the proportions of reactions that are reversible in each GSMM: 46% of all reactions in Human-GEM are reversible, 40% in yeast-GEM, and only 24% in iML1515. These differences may exist because more experiments to determine the extent to which particular metabolic reactions are reversible have been performed in *S. cerevisiae* and *E. coli* than in human cells, and the curators of Human-GEM may have erred on the side of making reactions reversible rather than irreversible in the absence of direct experimental evidence.

Another notable difference across organisms is that there appear to be significantly more dead-end reactions in glycan metabolism in yeast-GEM than either Human-GEM or iML1515 (Fig. 2). This is largely attributable to the fact that we only categorized 40 reactions in all of yeast-GEM as glycan metabolism reactions (see “Methods”), as opposed to 101 in iML1515 and 626 in Human-GEM. As glycan metabolism is generally considered to be a highly conserved part of metabolism [67], this discrepancy in the number of glycan metabolism reactions in each GSMM is more likely to be attributable to differences in how the three GSMMs were curated than biological differences in glycan metabolism between human, *S. cerevisiae*, and *E. coli* cells *in vivo*.

The dilution test also flags a higher proportion of reactions in yeast-GEM and iML1515 compared to Human-GEM (Fig. 2). This is especially true of reactions involving cofactors, glycans, lipids, and secondary metabolites in iML1515 and of reactions involving lipids and other amino acids in yeast-GEM. The reasons for this variability are currently unknown, and future investigation of the reactions flagged by the dilution test may determine whether these differences arise from specific curation errors localized in distinct regions of the GSMMs.

### MACAW uses network inference and visualization to help identify and streamline GSMM curation

A number of existing tools enumerate potential errors in GSMMs, but do relatively little to suggest what changes to the GSMM would correct the highlighted errors. The tests in MACAW attempt to provide additional information about each reaction that they highlight to suggest which changes would be necessary to fix the potential problem with the reaction. For example, for each reaction that the duplicate test flags, it also provides a list of other reactions in the GSMM that are likely duplicates so that one can compare all reactions in each group side-by-side to assess whether any should be removed, combined, or otherwise edited. Similarly, for each reaction flagged by the dead-end test, MACAW identifies which of the metabolites that participate in that reaction connect it to other dead-end reactions or are the “root” of the dead-end (i.e., a metabolite that can be produced but not consumed by all reactions it participates in or vice versa). This can help reveal cases where many dead-end reactions all connect back to a single dead-end metabolite, such that adding a single new reaction to the GSMM would resolve the issues with many reactions simultaneously.

Some groups of dead-end reactions cannot be traced back to any single dead-end metabolite, such as the short series of reactions involving alanyl-histidine in yeast-GEM (Additional File 1: Figure S1). None of these reactions are capable of sustaining steady-state fluxes, as alanyl-histidine can only be consumed in both the extracellular compartment (by r_4511) and the vacuolar compartment (by r_4515), and the only other reactions involving alanyl-histidine in yeast-GEM are only capable of transporting it between different compartments. Cases like these are particularly challenging to make sense of just by looking at a list of all dead-end reactions in a particular GSMM, so the dead-end test in MACAW produces both a list of all dead-end metabolites and a network connecting them. GSMMs are often challenging to visualize, as they generally contain thousands of densely interconnected reactions that produce uninterpretable hairballs if all reactions and metabolites are visualized simultaneously. These dead-end networks allow one to visualize each connected pathway of dead-end reactions as relatively small and interpretable subnetworks. Due to the way that the dead-end test constructs these networks (see “Methods”), each one likely represents relatively few underlying errors in the GSMM compared to the number of reactions in the subnetwork.

The other tests in MACAW also connect the reactions they flag into similar networks of related reactions, and MACAW also contains an algorithm for merging the networks produced by each test into a single comprehensive network. Using the three manually curated GSMMs from earlier (Human-GEM, yeast-GEM, and iML1515) as examples, each is dominated by a large connected component containing reactions flagged by many different tests surrounded by many much smaller connected components that each largely or exclusively consist of reactions flagged by the same test (Fig. 3A, Additional File 1: Figures S2A, S3A). In each case, the largest connected component contains the biomass reaction, whose reactants encompass a wide variety of metabolites from across each metabolic network. Reactions flagged by the loop and duplicate tests tend to be in smaller connected components than reactions flagged by the dilution and dead-end tests (Fig. 3B, Additional File 1: Figures S2B, S3B), which is generally consistent with how each test connects the reactions it flags into networks (see “Methods”). These merged networks can also help explain why some reactions were flagged by multiple different tests within MACAW (Fig. 3C), such as a pair of duplicate reactions that involve the same metabolites but are oriented in opposite directions (i.e., the products of one are the reactants of the other) that collectively form a loop capable of sustaining unlimited flux.

**Figure 3.**
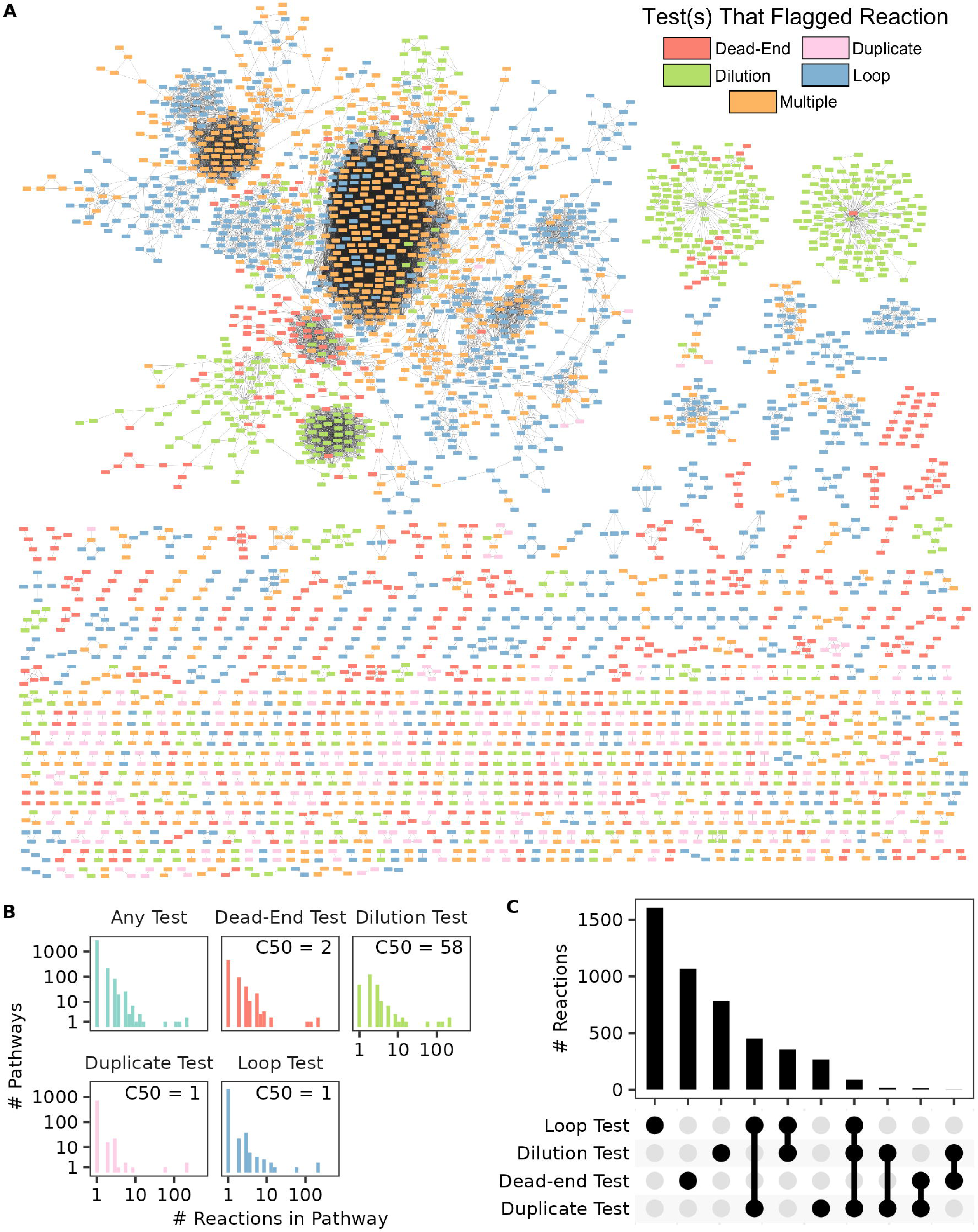
Overview of reactions in version 1.15 of Human-GEM flagged by one or more tests in MACAW. (**A**) Each node represents a single reaction; see “Methods” for explanation of how reactions were connected. The color of each node indicates which test(s) the reaction was flagged by. (**B**) Distributions of numbers of reactions in each connected component (“pathway”) shown in (**A**) for all pathways or only pathways containing at least one reaction flagged by the specified test. The C50 of each distribution is the median number of reactions flagged by each test weighted by the total number of reactions in the same pathway as each reaction. (**C**) UpSET plot showing number of reactions flagged by each observed combination of tests in MACAW.

Using the networks of flagged reactions generated by MACAW for version 1.15 of Human-GEM, we identified and proposed fixes for approximately 700 reactions, representing most of the corrections that were approved and incorporated into Human-GEM by its curators between versions 1.15 and 1.18. This is remarkable, considering that version 1.15 of Human-GEM already represented decades of curation efforts by dozens of other curators. Our curation efforts have resulted in a modest but noticeable reduction in the proportion of reactions flagged by at least one test over the course of the recent versions of Human-GEM (Additional File 1: Figure S4A). The reactions that were flagged by one or more tests in version 1.15 but not version 1.18 were not particularly concentrated in any one pathway (Additional File 1: Figure S4B). The largest difference between the two versions is in the proportion of energy metabolism reactions flagged by the loop test, which we discuss in greater detail below.

### Application of MACAW to automatically generated GSMMs

To more directly test whether differences in the proportions of reactions flagged by particular tests between GSMMs are more attributable to different curation approaches than genuine biological differences between the modeled organisms, we ran MACAW on a collection of 83 GSMMs for *Bordetella pertussis*, *Lactobacillus plantarum*, and *Pseudomonas putida* created using a variety of different methods for reconstructing GSMMs [57]. These reconstruction methods differ in the sources of data they draw on to reconstruct a GSMM (e.g., KEGG [66], MetaCyc [68], ModelSEED [69]), how they incorporate data into a GSMM (e.g., deciding which new reactions to add or which present reactions to remove), and the extent to which they use existing GSMMs as templates (as opposed to reconstructing a GSMM from an annotated genome *de novo*). In these GSMMs, the proportion of reactions flagged by each test appears to be more strongly influenced by the method used to create the GSMM than the organism each GSMM represents (Fig. 4A). The GSMMs produced by CarveMe [70] stand out for having exceptionally few reactions flagged by the dead-end test relative to the others. This is attributable to the fact that CarveMe begins with a “universal” GSMM representing most reactions known to be catalyzed by any prokaryote and prunes this GSMM, which has been manually curated to contain no dead-ends into one for the target organism. The other reconstruction methods either construct GSMMs by editing existing GSMMs in databases such as BiGG that generally contain numerous dead-ends and/or by adding individual new reactions from databases of reactions without necessarily ensuring the added reactions do not wind up as dead-ends. It is worth noting that CarveMe is only capable of reconstructing GSMMs for prokaryotes, while the other methods can be applied to any organism. Previous work has suggested that CarveMe produces models of comparable quality to manually curated models. Indeed, we observed that the overall proportions of reactions flagged by any test in the GSMMs produced by CarveMe (33%-36%) are similar to the proportions in the three manually curated models mentioned earlier (Human-GEM, yeast-GEM, and iML1515; 36%-52%).

**Figure 4.**
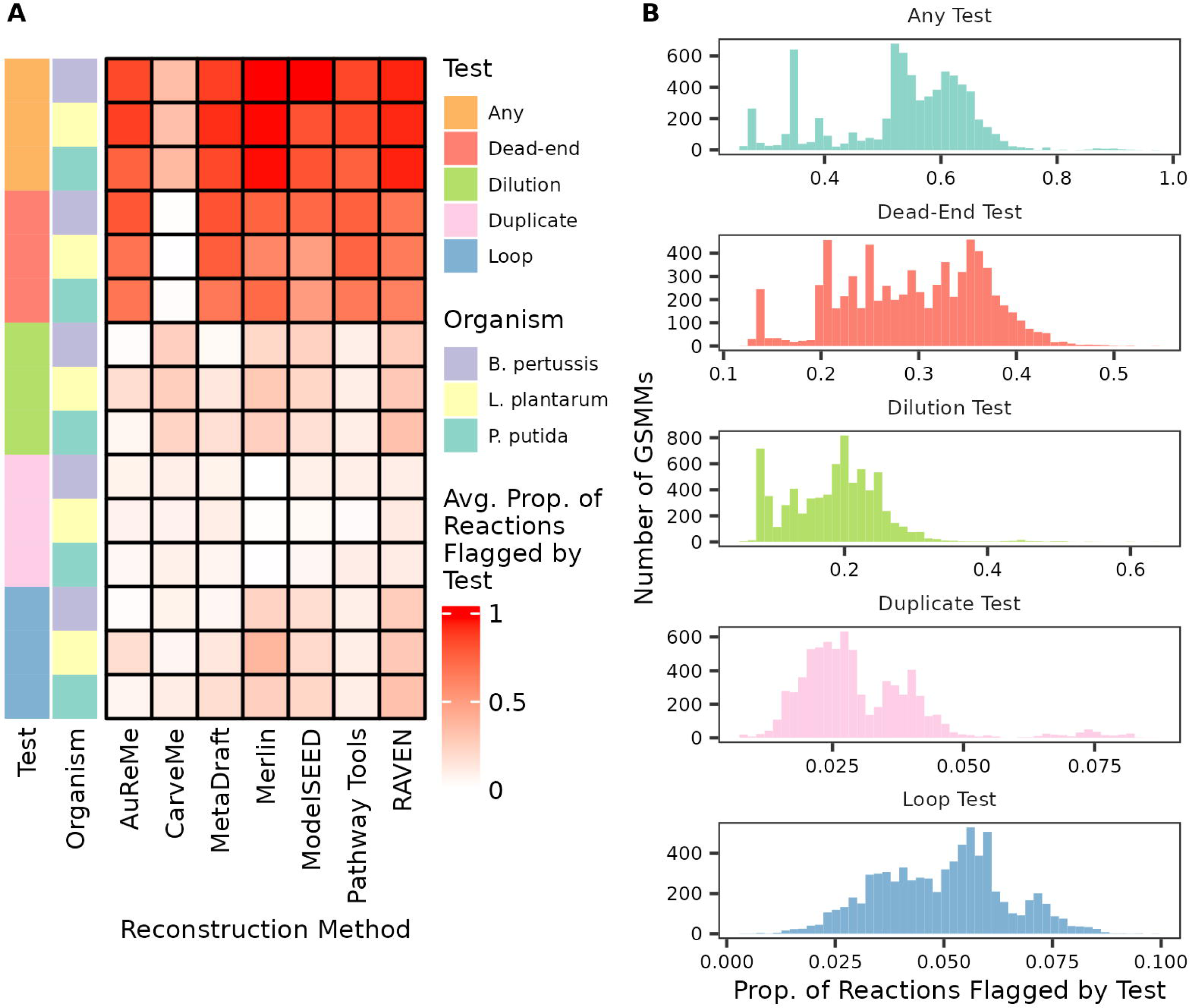
Proportions of reactions flagged by each test in automatically curated GSMMs. (**A**) Average proportion of reactions flagged by each test in the GSMMs made with the indicated reconstruction method for the indicated organisms. (**B**) Distributions of proportions of reactions flagged by each test in each of the AGORA2 GSMMs.

Our previous analysis of GSMMs for three bacteria suggested that the proportion of reactions flagged by each test in MACAW are similar across GSMMs for organisms. To evaluate this more broadly, we applied MACAW to 7,302 GSMMs representing human gut microbes in the AGORA2 collection [56] (Fig. 4B). There is significant variability in the proportions of reactions flagged by each test across all of the AGORA2 GSMMs, and since all of the AGORA2 GSMMs were created using the same custom reconstruction pipeline, these differences are not attributable to differences in how each GSMM was created. Some of the distributions appear to be multimodal, especially the distribution of proportions of reactions flagged by the loop and duplicate tests, suggesting that certain subsets of the AGORA2 strains possess some quality that makes it challenging to avoid certain kinds of errors when constructing GSMMs for those strains.

### Using MACAW to resolve duplicate energy metabolism reactions in Human-GEM

To illustrate the applicability of MACAW for error correction, we highlight a few interesting subsets of reactions that were flagged by MACAW in version 1.15 of Human-GEM which lead us to discover and propose solutions for a number of errors, most of which have been incorporated into version 1.18. The majority of these reactions are redox reactions and were flagged by the duplicate test. Enzymes that catalyze redox reactions frequently use cofactors such as NAD(H), FAD(H_2_), and ubiquinone. We found many pairs of redox reactions in Human-GEM that only differed by the identity of the cofactor, which do not always represent errors in a GSMM, but we have found that many such pairs misrepresent the corresponding reaction in the target organism. For example, Succinate Dehydrogenase (SDH) can oxidize succinate to fumarate while reducing its FAD cofactor, then use the FADH_2_ to reduce ubiquinone to ubiquinol [71]. Version 1.15 of Human-GEM represented this reaction twice: both as a series of two reactions, one between succinate/fumarate and FAD(H_2_) and the other between FAD(H_2_) and ubiquinone/ubiquinol (Fig. 5A, left) and as a single reaction that directly transfers electrons between succinate/fumarate and ubiquinone/ubiquinol (Fig. 5A, right). We chose to remove the FAD-dependent reaction (Fig. 5A, leftmost) to reflect the fact that, unlike most human FAD-dependent enzymes, the FAD cofactor in SDH is bound covalently, so the FAD bound to SDH is sequestered from the pool of non-covalently-bound FAD that is effectively shared by most other human FAD-dependent enzymes [72–74]. In addition to resolving this duplicate/redundant representation of a single enzyme activity, this change also removed the risk of predicting a thermodynamically infeasible loop flux between succinate, fumarate, FAD(H_2_), ubiquinone, and ubiquinol using these three reactions. While a number of techniques exist for dealing with such loop fluxes when using GSMMs to predict fluxes [38,62,63,75–77], they are often challenging to use and/or depend on questionable assumptions about biology/biochemistry, and removing this one reaction is a relatively simple way to resolve the issue in this instance.

**Figure 5.**
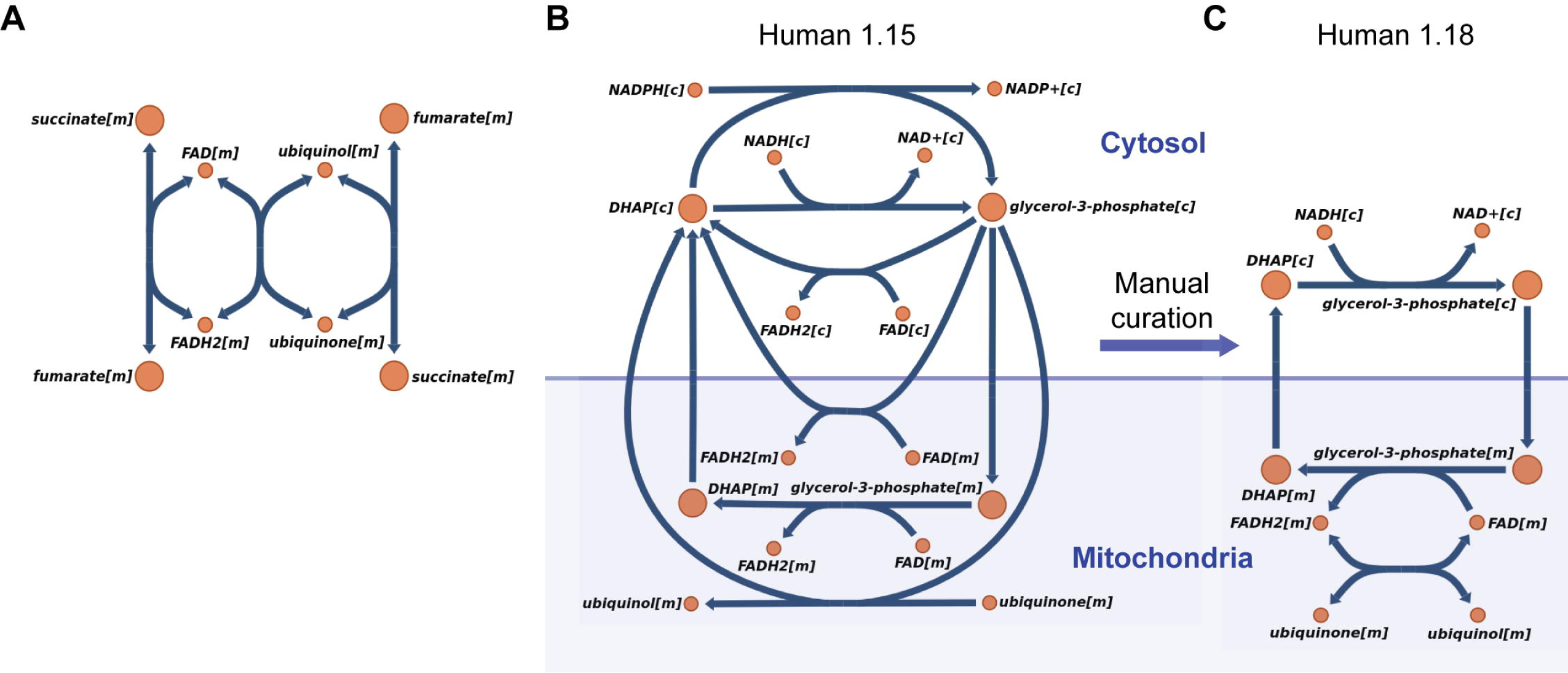
Duplicate redox reactions in Human-GEM. (**A**) Escher map of the reactions representing the activity of succinate dehydrogenase in version 1.15 of Human-GEM. Letters in square brackets indicate the subcellular compartment of each metabolite: “c” for cytoplasmic metabolites and “m” for mitochondrial. (**B**) Escher map of the reactions representing the glycerol-3-phosphate shuttle in version 1.15 of Human-GEM. (**C**) Escher map of the reactions representing the glycerol-3-phosphate shuttle after manual curation in version 1.18 of Human-GEM.

The duplicate test also revealed similar but slightly more complicated problems with the representation of the Glycerol-3-Phosphate (G3P) shuttle (Fig. 5B). The first step of the G3P shuttle is the reduction of DihydroxyAcetone Phosphate (DHAP) to G3P by either GPD1 or GPD1L in the cytosol. GPD1 and GPD1L both use NADH as an electron donor [78,79], yet version 1.15 of Human-GEM also contained a version of this reaction that incorrectly used NADPH (Fig. 5B). G3P produced by GPD1 or GPD1L is transported into the mitochondrial intermembrane space, where GPD2 oxidizes it back into DHAP. While this DHAP is ultimately transported back into the cytosol, version 1.15 of Human-GEM had two versions of the reaction catalyzed by GPD2 that involved cytosolic G3P and DHAP in addition to two versions that used mitochondrial G3P and DHAP (Fig.5B). Like SDH, GPD2 also uses an FAD cofactor as an intermediate electron carrier, first reducing it while oxidizing G3P, then oxidizing it while reducing ubiquinone in the inner mitochondrial membrane [80]. Correcting these errors significantly simplified the representation of the G3P shuttle in version 1.18 of Human-GEM (Fig. 5C). While this set of duplicates did not create a loop like the duplicate SDH reactions did (Fig. 5A), their presence made it difficult to interpret predicted fluxes through the G3P shuttle, a central component of energy metabolism in human cells.

### Using MACAW to improve knockout prediction by correcting errors with lipoic acid metabolism in Human-GEM

The dilution test highlighted most of the reactions that represent lipoic acid metabolism in version 1.15 of Human-GEM, including the Glycine Cleavage System (GCS), and Pyruvate Dehydrogenase (PDH) (Fig. 6A). The metabolite produced by the lipoic acid biosynthesis reaction (lipoyl-GCSH) cannot be converted into the metabolite that represents lipoic acid’s role as a cofactor in the GCSH and PDH reactions (lipoamide), so lipoamide can only be recycled between its oxidized, reduced (dihydrolipoamide), and conjugated (S-acetyldihydrolipoamide, S-aminomethyldihydrolipoamide) forms. Adding dilution constraints for any of these lipoic acid metabolites requires the total flux producing it to exceed the total flux through non-dilution reactions consuming it, which is impossible without a connection between the biosynthesis reaction and the GCS and PDH reactions.

**Figure 6.**
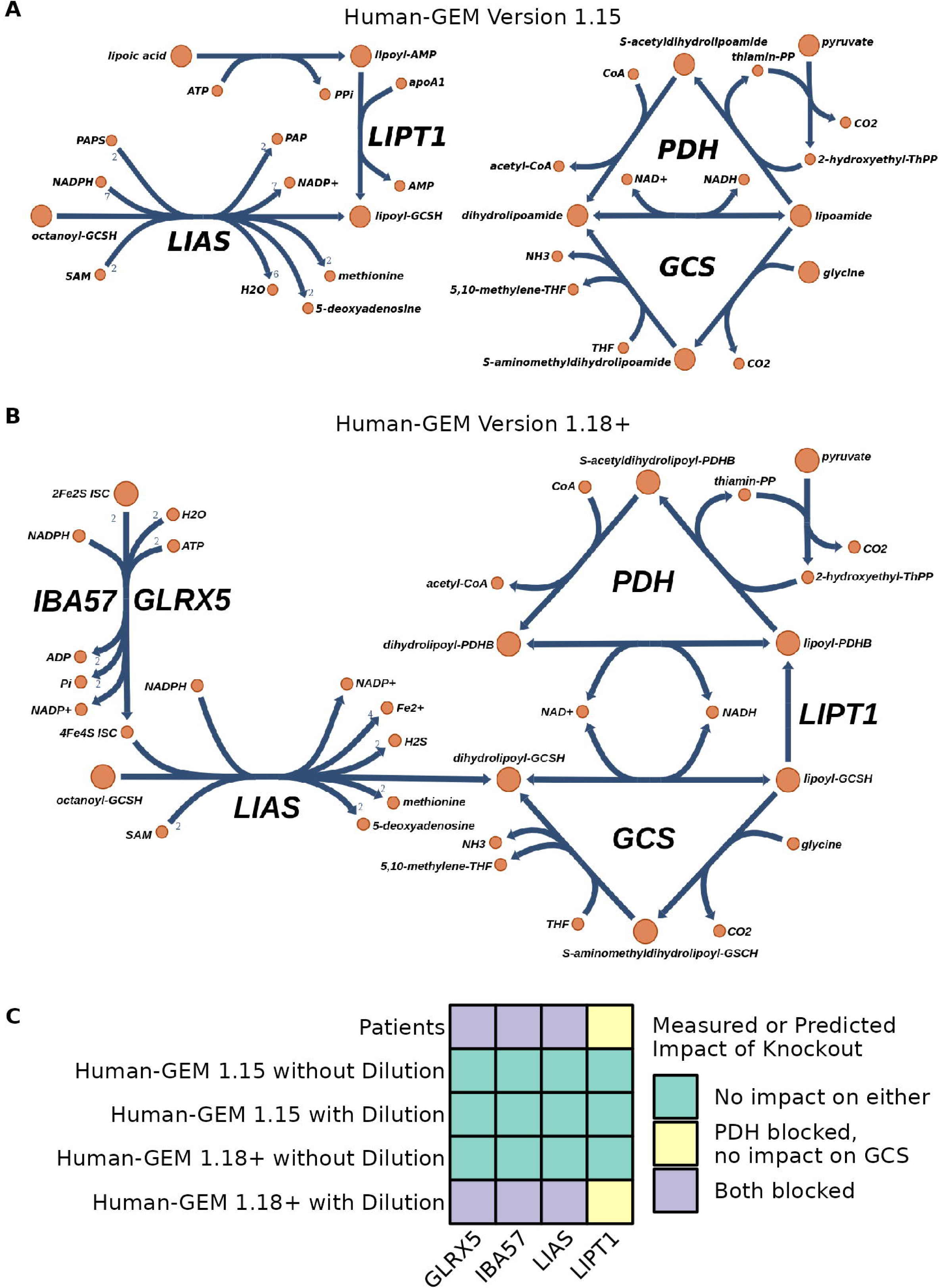
Correcting errors in lipoic acid metabolism in Human-GEM. (**A**) Escher map of the reactions in version 1.15 of Human-GEM that represent the enzymatic activities of Lipoic Acid Synthetase (LIAS), Lipoyl Transferase 1 (LIPT1), Glycine Cleavage System Protein H (GCSH), and Pyruvate Dehydrogenase E1 Subunit Beta (PDHB). Protons have been omitted for clarity. The metabolite labeled “octanoyl-GCSH” is named “[protein]-N6-(octanoyl)lysine” in version 1.15 Human-GEM, and “lipoyl-GCSH” is “[protein]-N6-(lipoyl)lysine.” (**B**) Escher map of the reactions representing the enzymatic activities of all of the enzymes shown in (**A**) as well as Iron-Sulfur Cluster Assembly Factor IBA57 (IBA57) and Glutaredoxin 5 (GLRX5) after correcting the errors in lipoic acid metabolism that were present in version 1.18 of Human-GEM. (**C**) Heatmap comparing the observed impacts of loss-of-function mutations in GLRX5, IBA57, LIAS, and LIPT1 on fluxes through the Glycine Cleavage System (GCS) and Pyruvate Dehydrogenase (PDH) in human patients to the predicted impacts of knocking out each gene on fluxes through the corresponding reactions in different versions of Human-GEM with or without dilution constraints imposed. Version 1.18 of Human-GEM only corrects some of the errors with lipoic acid metabolism that were present in version 1.15, so Human-GEM 1.18+ corresponds to a version of Human-GEM 1.18 with all remaining problems fixed, such that the pathway looks like panel (**B**).

Correcting this issue is more complicated than simply adding a new reaction to convert lipoyl-GCSH into lipoamide (or combining the two into a single metabolite), as the two metabolites are also in different subcellular compartments of version 1.15 of Human-GEM. Lipoyl-GCSH only exists in the cytosolic compartment, while lipoamide and its derivatives only exist in the mitochondrial compartment. Lipoic acid biosynthesis occurs exclusively in the mitochondria of human cells, as do all the enzymes that use lipoic acid as a cofactor [81].

Moving the biosynthesis reaction to the mitochondrial compartment is complicated by the fact that several of its metabolites only exist in the cytosolic compartment of version 1.15 of Human-GEM. One such metabolite is 3′-phosphoadenosine-5′-phosphosulfate (PAPS), which should not participate in this reaction: the sulfur atoms in lipoic acid are derived from one of the iron-sulfur clusters associated with LIAS as opposed to PAPS [82,83]. Version 1.15 of Human-GEM also completely lacked the iron-sulfur cluster biosynthesis pathway, so correcting the mislocalization of the lipoic acid biosynthesis reaction also involved changing several of its reactants and products and adding in a number of other missing reactions (Fig. 6B). In addition to misrepresenting the subcellular localization of lipoic acid biosynthesis, version 1.15 of Human-GEM also misrepresented the role of LIPT1. While LIPT1 was previously thought to be capable of attaching free lipoic acid to GCSH, it is now known to only be capable of transferring lipoic acid that is bound to GCSH (LIAS converts an octanoyl moiety bound to GCSH into a lipoyl moiety that remains bound to GCSH [81]) to a number of other enzymes, including the beta subunit of the PDH complex (Fig 6B) [84].

The misrepresentation of LIPT1 and the absence of the iron-sulfur cluster biosynthesis pathway rendered version 1.15 of Human-GEM incapable of accurately predicting the impacts of knockouts in several genes associated with these pathways. Patients with mutations in GLRX5 or IBA57 (two of the enzymes involved in iron-sulfur cluster biosynthesis [85,86]) or LIAS frequently have elevated levels of lactate and glycine in their blood, which is thought to be caused by low activity of the GCS and PDH [81]. Patients with mutations in LIPT1 also have elevated levels of lactate but have normal levels of glycine [81]. Due to the errors described above, version 1.15 of Human-GEM fails to accurately predict the consequences of these mutations (Fig. 6C).

Correcting all of the errors we identified in Human-GEM allows it to accurately predict the impacts of knockouts in all four genes (Fig. 6C). As version 1.18 of Human-GEM (the most recent at the time of writing) only corrects some of the above issues with lipoic acid metabolism, we refer to the version of Human-GEM in which we corrected all of the above errors in lipoic acid metabolism as version 1.18+. Note that the predicted knockouts only align with the observed phenotypes of individuals with mutations in these genes when dilution constraints are imposed on version 1.18+ of Human-GEM. While the present work primarily focuses on dilution constraints as tools for identifying errors in GSMMs, previous applications of dilution constraints have primarily discussed their capacity to improve the accuracy of knockout predictions [60,61].

## Discussion

We have created MACAW, an ensemble of network-based algorithms for identifying and visualizing potential errors in GSMMs. MACAW incorporates variations of previously published approaches for highlighting errors with individual reactions and extends them to also identify errors which only become apparent at the level of whole pathways of reactions. By following up on errors highlighted by MACAW in Human-GEM, we have demonstrably improved its capacity to accurately predict the impacts of knockouts. More generally, we have also used MACAW to analyze how different approaches to reconstruction and curation of GSMMs affect the kinds of errors they contain.

MEMOTE [51] and ErrorTracer [52] resemble MACAW in that they are also collections of algorithms for highlighting potential errors in arbitrary GSMMs, but differ in a number of key respects (Table 1). While some of the tests that comprise MACAW are similar to tests included in MEMOTE and ErrorTracer, neither of them joins the reactions flagged by their tests into networks in the way that MACAW does. ErrorTracer connects all reactions in the given GSMM into a single network, while MACAW only joins reactions flagged by at least one test. This is particularly helpful in the case of the loop test, which has equivalents in both MEMOTE and ErrorTracer, but both tools only report which reactions were flagged by the loop test, with no further information. MACAW also groups the reactions flagged by the loop test into distinct loops, which significantly simplifies the process of understanding what changes to the component reactions would be necessary to resolve the loop. The duplicate test in MEMOTE only highlights pairs of reactions that are exactly identical in every way, while the duplicate test in MACAW also highlights pairs of reactions that are largely identical but have different stoichiometric coefficients, go in different directions, etc. The overwhelming majority of potential duplicates flagged by MACAW in every GSMM we have tested it on so far are not exact duplicates, suggesting that the number of errors that MEMOTE fails to identify by solely focusing on exact duplicates is considerable. However, it is important to note that the goal of the duplicate test in MACAW is to highlight groups of reactions that are worth examining closely for potential duplicates, as opposed to highlighting reactions that definitely represent an error in the GSMM. MEMOTE also contains many tests unrelated to those in MACAW, many of which focus on very fundamental errors in GSMMs that would prevent them from being used for most potential applications of GSMMs. The two tools are more complementary than redundant: the tests in MEMOTE serve as tools for getting GSMMs to a basic level of functionality and usability, while the tests in MACAW facilitate further refinement of GSMMs beyond that point.

The large proportions of reactions found to be dead-ends in most of the GSMMs tested, especially the manually curated ones (Human-GEM, yeast-GEM, and iML1515), should not necessarily be interpreted as a metric of their quality. While GSMMs are commonly used to predict steady-state metabolic fluxes, for which dead-end reactions are useless, GSMMs are also used as repositories of biochemical knowledge about a particular organism. As our knowledge of the metabolic capacities of many organisms is largely incomplete [87], a GSMM that contains dead-end reactions may simply accurately reflect the current state of our understanding. A GSMM that contains no dead-end reactions may seem more useful or complete in the context of predicting fluxes, but might fail to represent dozens or hundreds of reactions known to occur in the modeled organism, and thus be less useful as a repository of knowledge. The proportion of dead-end reactions in a given GSMM may have more to say about how much remains to be discovered about the organism it represents than the accuracy of the GSMM. The dead-end test can highlight opportunities for new experiments to fill gaps in the current understanding of a particular organism’s metabolic capabilities.

The dilution constraints used in the dilution test are inspired by the dilution constraints used in previous works to alter predicted fluxes and the impacts that knockouts have on those predicted fluxes [60,61]. This is, to our knowledge, the first application of dilution constraints to the context of highlighting errors in GSMMs during the curation process. While previous works have interpreted the fluxes through dilution reactions as representing the dilution of intermediate metabolites as cells grow and divide, we believe it is also interesting to consider them as representing loss of intermediate metabolites to side reactions, such as with reactive oxygen species. This also relates to our choice to constrain the fluxes through dilution reactions to be some fraction of the sum of the non-dilution fluxes involving the metabolite being diluted: all chemical reactions involve the formation of unstable transition states that are frequently capable of forming into more than just two sets of reactants or products, each with varying probabilities of occurring [88,89]. The more times a particular transition state forms, the more times it will resolve into less common sets of products, thus the dilution flux for a given metabolite scales with the total magnitude of the non-dilution fluxes involving that metabolite. While we identified several errors in lipoic acid metabolism in Human-GEM using the dilution test, the corrected version of Human-GEM was only able to accurately predict the impact of knockouts in that pathway when dilution constraints were imposed for all metabolites (Fig. 6C). Dilution constraints can therefore play a multifaceted role in improving the quality of GSMMs.

A variety of tools exist for mitigating the problem of thermodynamically infeasible loops when using GSMMs to predict steady-state fluxes [38,62,63,75–77]. While they all certainly produce more realistic predictions than the infinite fluxes such loops are capable of sustaining otherwise, some are computationally expensive and thus difficult to apply to larger GSMMs, some require thermodynamic and/or kinetic data that is not known for most reactions in most organisms and is difficult to estimate accurately, and the less computationally and data-intensive approaches rely on approximations and heuristics that render the predictions questionably relevant in many biological contexts. Furthermore, all but the most trivial GSMMs (e.g., those that only represent a few “core” pathways like central carbon metabolism) have multiple configurations of fluxes through the network that satisfy all constraints [90], and few of the existing methods for mitigating loop fluxes are compatible with algorithms for characterizing distributions of all possible fluxes through a GSMM [91–93]. While loops are not always caused by errors in the construction of a GSMM, many are. Identifying such loops using MACAW’s loop test can allow one to mitigate this problem before engaging with the fraught question of how best to predict fluxes from a GSMM.

### Conclusions

We present MACAW, a tool for identifying and guiding the correction of errors in GSMMs. Our approach is particularly suited for identifying and correcting subtle errors in GSMMs that only become apparent when considering multiple reactions collectively. MACAW identifies numerous errors in both manually curated and automatically generated GSMMs, highlighting opportunities for improving current approaches to creating and refining GSMMs. Overall, MACAW represents a useful and innovative addition to the set of tools available for improving the quality of GSMMs, both in their capacity as repositories of biochemical knowledge and in their capacity as tools for predicting metabolic phenotypes.

## Methods

### Genome-Scale Metabolic Models

All versions of Human-GEM [53] were downloaded from GitHub (https://github.com/SysBioChalmers/Human-GEM), as was version 9.0.0 of yeast-GEM [54] (https://github.com/SysBioChalmers/yeast-GEM) and the assortment of GSMMs for *B. pertussis*, *L. plantarum*, and *P. putida* (https://github.com/SystemsBioinformatics/pub-data/tree/master/reconstruction-tools-assessment). iML1515 [55] was downloaded from BiGG [94], and the AGORA2 collection was downloaded from the Virtual Metabolic Human database [95].

Before running MACAW on all versions of Human-GEM, lower bounds on the fluxes through all exchange reactions (representing the maximum allowed uptake of each metabolite from the external environment) were set to −1,000 for all metabolites present in DMEM or fetal bovine serum (FBS) (Table S1) and 0 for all others. We similarly constrained the exchange reactions in yeast-GEM using the composition of a minimal mineral medium commonly used in isotope tracing experiments [96] (Table S2) and exchange reactions in iML1515 using the composition of M9 media [97] (Table S3). Bounds on exchange reactions were left as-is for all of the GSMMs for all of the GSMMs shown in Fig. 4.

### The Dead-End Test

The dead-end test checks every metabolite in the given GSMM, one at a time, to see if it can only be consumed by every reaction it participates in or only be produced by every reaction it participates in. Every time it finds a metabolite meeting either criterion, it adds that metabolite to the list of dead-end metabolites and all reactions it participates in to the list of dead-end reactions. Each time a new metabolite is added to the list of dead-end metabolites, this test also re-evaluates all other metabolites that participate in any of the reactions that the newly identified dead-end metabolite participates in, unless they are already in the list of dead-end metabolites. When a metabolite participates in exactly one reversible reaction and at least one irreversible reaction, if the metabolite is a product in all of the irreversible reactions or a reactant in all of the irreversible reactions, the one reversible reaction is made irreversible in the appropriate direction (e.g., if one of its products can only be consumed by every other reaction it participates in, it is made irreversible in its forward direction by setting the lower bound on its flux to 0), and all other metabolites that participate in the reversible reaction are retested.

Once all metabolites have been checked at least once, the dead-end test generates a table indicating which reactions were found to be dead-ends, all dead-end metabolites that participate in each flagged reaction, and which reversible reactions had metabolites that could only be produced or only be consumed by every other reaction they participated in. It also creates an edge list describing a bipartite network in which nodes represent either dead-end reactions or dead-end metabolites, where all dead-end metabolites are connected to the dead-end reactions that they participate in.

### The Dilution Test

We define a “dilution” reaction for each metabolite that consumes it and produces nothing whose flux is constrained to be some fraction of the sum of the absolute values of the fluxes of all other reactions consuming or producing the metabolite in question. In a GSMM with i metabolites that participate in j reactions represented by a stoichiometric matrix S, the “dilution” flux for metabolite i is

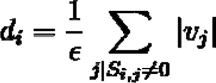

where v_j_ is the flux through reaction j, and ε is a free parameter that governs how large the flux through the “dilution” reaction is relative to the fluxes through the other reactions that metabolite i participates in. All results shown in this paper were obtained with ε = 1000 for all metabolites in each GSMM, but in principle these can be different values for different metabolites. Since the dilution test only flags reactions that are capable of zero flux after imposing a dilution constraint, the exact value of ε used does not impact which reactions are flagged (but may be more impactful in other applications of dilution constraints).

The “dilution constraints” require that every metabolite that participates in any reactions with non-zero fluxes to be capable of having the total flux producing that metabolite exceed the total flux consuming it, which prevents metabolites from being “perfectly” recycled via internal loops with no input from any external sources (i.e. exchange reactions). The dilution test adds a dilution reaction and constraint for each metabolite in the given GSMM one at a time, performs FVA for all reactions involving that metabolite, and records whether or not that dilution constraint prevented all reactions involving that metabolite from sustaining non-zero fluxes. It then uses the same algorithm as the dead-end test to determine whether any additional reactions beyond those that directly consume or produce each “dilution-blocked” metabolite would also be blocked by dilution constraints. The dilution test produces a list of all reactions found to be blocked by adding a dilution constraint for a single metabolite and an edge list describing a bipartite network of metabolites and reactions in the same fashion as the dead-end test.

The dilution test performs a number of pre-processing steps before imposing dilution constraints. First, it uses the results of the dead-end test on the same GSMM to set all bounds on dead-end reactions to zero in order to prevent a dilution reaction for a dead-end metabolite from allowing the other reaction(s) involving that metabolite to become capable of sustaining flux. Since significant proportions of reactions in most GSMMs are dead-end reactions, this step shortens the time required to run the dilution test by a considerable amount in most cases.

Then, it performs FVA on all reversible reactions and sets their upper and lower bounds to their maximum and minimum possible fluxes. Finally, it adds what we term “leakage” reactions that allow every metabolite that exists in multiple compartments in the GSMM (e.g., extracellular, cytosolic, mitochondrial) to move between those compartments with a flux of up to 1 in either direction. Without leakage reactions, the dilution test would highlight certain antiporter reactions that do not represent recycling loops of metabolites that lack biosynthesis pathways (Fig. S5), which would constitute “false positives”. By default, MACAW constrains the fluxes through leakage reactions to be no more than 1 in either direction, but in principle any value that is relatively large compared to the 1/ε term in the dilution constraints (1/1000 by default) is sufficient to ensure that the dilution test does not generate “false positives”.

### The Duplicate Test

All pairs of reactions where both reactions involve exactly the same set of metabolites are labeled as duplicates. They are further subdivided into “exact”, “stoichiometric”, or “directional” duplicates based on the extent of further similarities between the reactions. “Exact” duplicates have the same stoichiometric coefficients, proceed in the same direction (i.e., the products of one are also the products of the other), and are either both reversible or both irreversible. “Stoichiometric” duplicates have different stoichiometric coefficients for their identical sets of metabolites (e.g., A + B -> C vs 2A + B -> 2C). “Directional” duplicates proceed in different directions (i.e., the products of one are the reactants of the other) and/or consist of at least one irreversible reaction and at least one reversible reaction. All sets of reactions identified as any kind of duplicates may or may not be associated with the same genes. The duplicate test also produces an edge list describing a network in which each node represents a reaction and all reactions that were labeled as duplicates of each other are joined by an edge.

If a list of pairs of metabolite IDs that correspond to the oxidized and reduced forms of electron carrier metabolites (e.g. NAD(H), FAD(H_2_), ubiquinone/ubiquinol) and a list of metabolite IDs that correspond to protons (H^+^) are provided, the test will also identify sets of “redox” duplicates. Redox duplicates are groups of reactions that involve exactly the same metabolites aside from the provided electron carriers and protons, both convert the oxidized forms of one of the given pairs into the reduced forms (or vice versa), and do not exclusively involve electron carriers and protons. An example pair of redox duplicates would be the two representations of the reaction catalyzed by succinate dehydrogenase in version 1.15 of Human-GEM discussed above: succinate + FAD <-> fumarate + FADH_2_ and succinate + ubiquinone <-> fumarate + ubiquinol (Fig. 5A).

### The Loop Test

The loop test sets the upper and lower bounds on all exchange reactions in the given GSMM to 0, removes any objective functions present (e.g., maximizing biomass production), then performs FVA and labels all reactions that are capable of non-zero fluxes as “in loops”. Then it generates 1,000 possible solutions to the GSMM using Cobrapy’s built-in implementation of OptGP [91], gets pairwise correlations between the distributions of possible fluxes for all reactions, and generates an edge list connecting all pairs of reactions with a pairwise correlation whose absolute value is above a certain threshold (0.9 by default) that also share at least one metabolite.

### Merging Test-Specific Edge Lists

Each test in MACAW produces a list of all reactions flagged by that test and an edge list defining a network connecting those reactions. MACAW also contains an algorithm for merging the networks produced by the different tests into a single comprehensive network. First, all metabolites present in the bipartite networks generated by the dead-end test and dilution test are connected to all reactions that they participate in that were flagged by the duplicate or loop tests. Then, a new edge is created between all pairs of reactions that are both connected to the same metabolite (note that this does not connect all reactions that share *any* metabolites; only reactions that share metabolites that were flagged by the dead-end or dilution tests). Finally, all nodes representing metabolites are removed, creating a monopartite network in which all nodes represent reactions.

### Categorizing Reactions Using The KEGG Orthology Database

KEGG organizes genes, proteins, and enzymes into a hierarchy of “functional orthologs”, each of which represents a group of genes/proteins/enzymes from different organisms that share similar functions and sequences [66,98]. All reactions in Human-GEM, yeast-GEM, and iML1515 were associated with KEGG functional orthologs via the gene IDs associated with each reaction; reactions that had no associated genes were labeled as “Not in KEGG”. Exactly one reaction out of all reactions in all three GSMMs was in the “Not included in regular maps” group, so that reaction was labeled as “Not in KEGG” for purposes of Fig. 2 and Additional File 1: Figure S4B. The labels that appear on the x-axes of Fig. 2 and Additional File 1: Figure S4B were derived from the labels of the largest groups in the hierarchy of KEGG functional orthologs in the “metabolism” branch.

### Data Analysis and Visualization

All data was generated and analyzed in Python and R [99,100]. Parts of Fig. 3 and Additional File 1: Fig. S2 and S3 were made in Cytoscape [101], and parts of Fig. 5 and 6 and Additional File 1: Fig. S1 and S5 were made with Escher [102]. All other data visualization was done in R [100,103–111]. All scripts used to generate, process, and visualize data are available on GitHub (https://github.com/Devlin-Moyer/macaw).

## Declarations

### Ethics approval and consent to participate

Not applicable.

## Consent for publication

Not applicable.

## Availability of data and materials

MACAW and all data generated and/or analyzed are available on GitHub at https://github.com/Devlin-Moyer/macaw [112].

## Authors’ contributions

DM, DS, and JIFB conceptualized the overall project and goals. DM wrote MACAW and all scripts used to generate data and figures. DM, DS, and JIFB discussed all analyses and results. DM and JR investigated reactions highlighted by one or more tests in Human-GEM, determined if there was an underlying error, and determined how to modify Human-GEM to correct the errors. DM, DS, and JIFB wrote the first draft of the manuscript, and all authors edited, proofread, and approved the final manuscript.

## Acknowledgements

We are grateful to Drs. Feiran Li, Hao Wang, and Jonathan Robinson, who helped determine which changes to make to Human-GEM to correct the errors highlighted by MACAW; and to Helen Scott, Michael Silverstein, and the other members of the Fuxman Bass and Segrè labs for feedback on the figures and manuscript.

## Supplemental information

Additional File 1.pdf Figures S1-S5.

Table S1.csv List of Human-GEM metabolite IDs for all metabolites present in Dulbecco’s Modified Eagle Medium and fetal bovine serum.

Table S2.csv List of yeast-GEM metabolite IDs for all metabolites present in a minimal mineral medium for *S. cerevisiae* [96].

Table S3.csv List of iML1515 metabolite IDs for all metabolites present in M9 minimal medium.

