## Supplementary figures and tables for "Semi-Automatic Detection of Errors in Genome-Scale Metabolic Models"

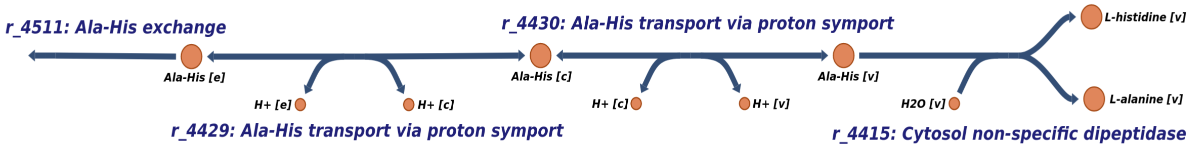


**Supplementary Figure 1. Escher map of a pathway of reactions flagged by the dead-end test in version 9.0.0 of yeast-GEM.**

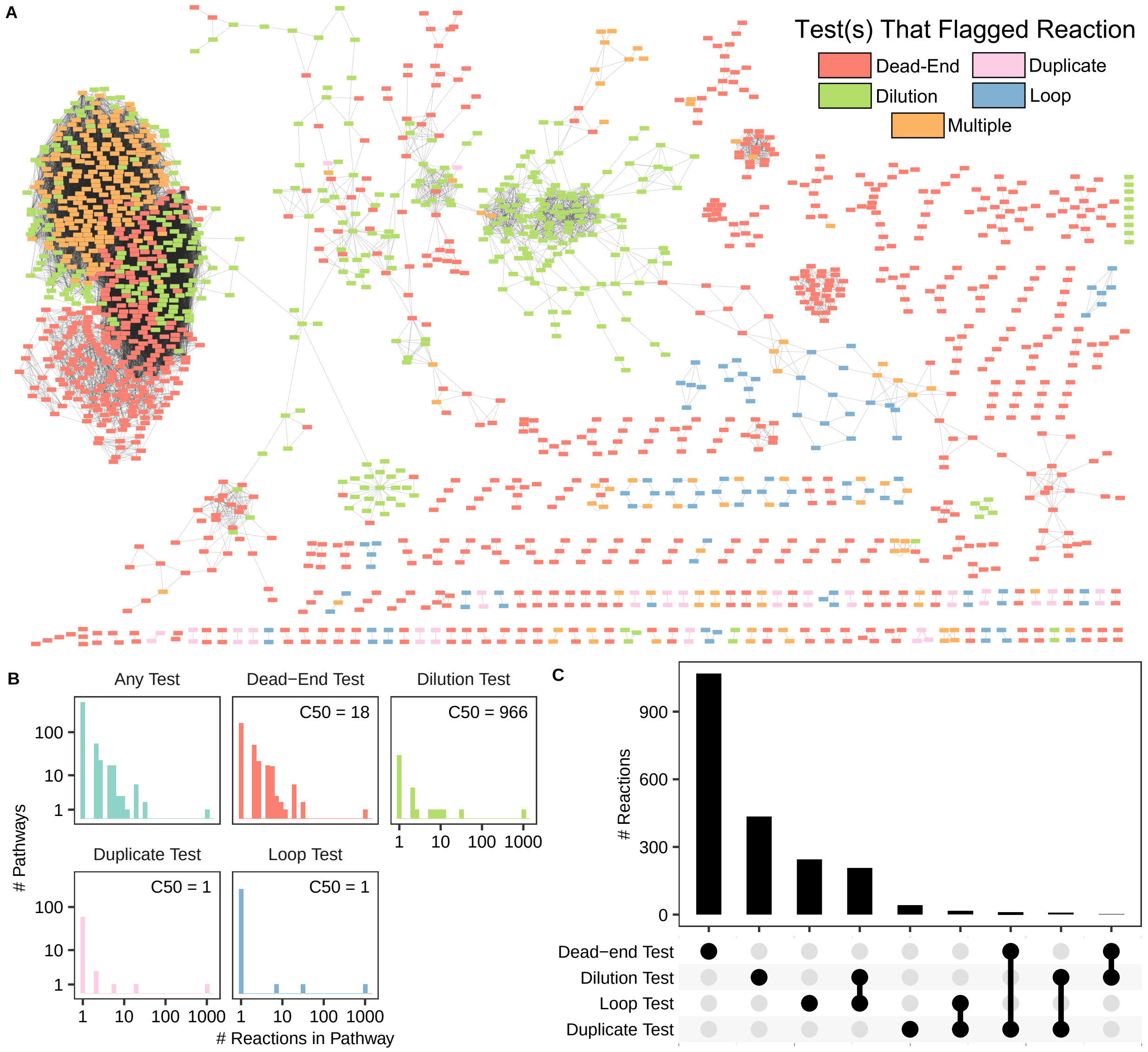


**Supplementary Figure 2. Overview of reactions in version 9.0.0 of yeast-GEM flagged by one or more tests in MACAW.** (A) Each node represents a single reaction; see Methods for explanation of how reactions were connected. The color of each node indicates which test(s) the reaction was flagged by. (B) Distributions of numbers of reactions in each connected component ("pathway") shown in (A) for all pathways or only pathways containing at least one reaction flagged by the specified test. The C50 of each distribution is the median number of reactions flagged by each test weighted by the total number of reactions in the same pathway as each reaction. (C) UpSET plot showing number of reactions flagged by each observed combination of tests in MACAW.

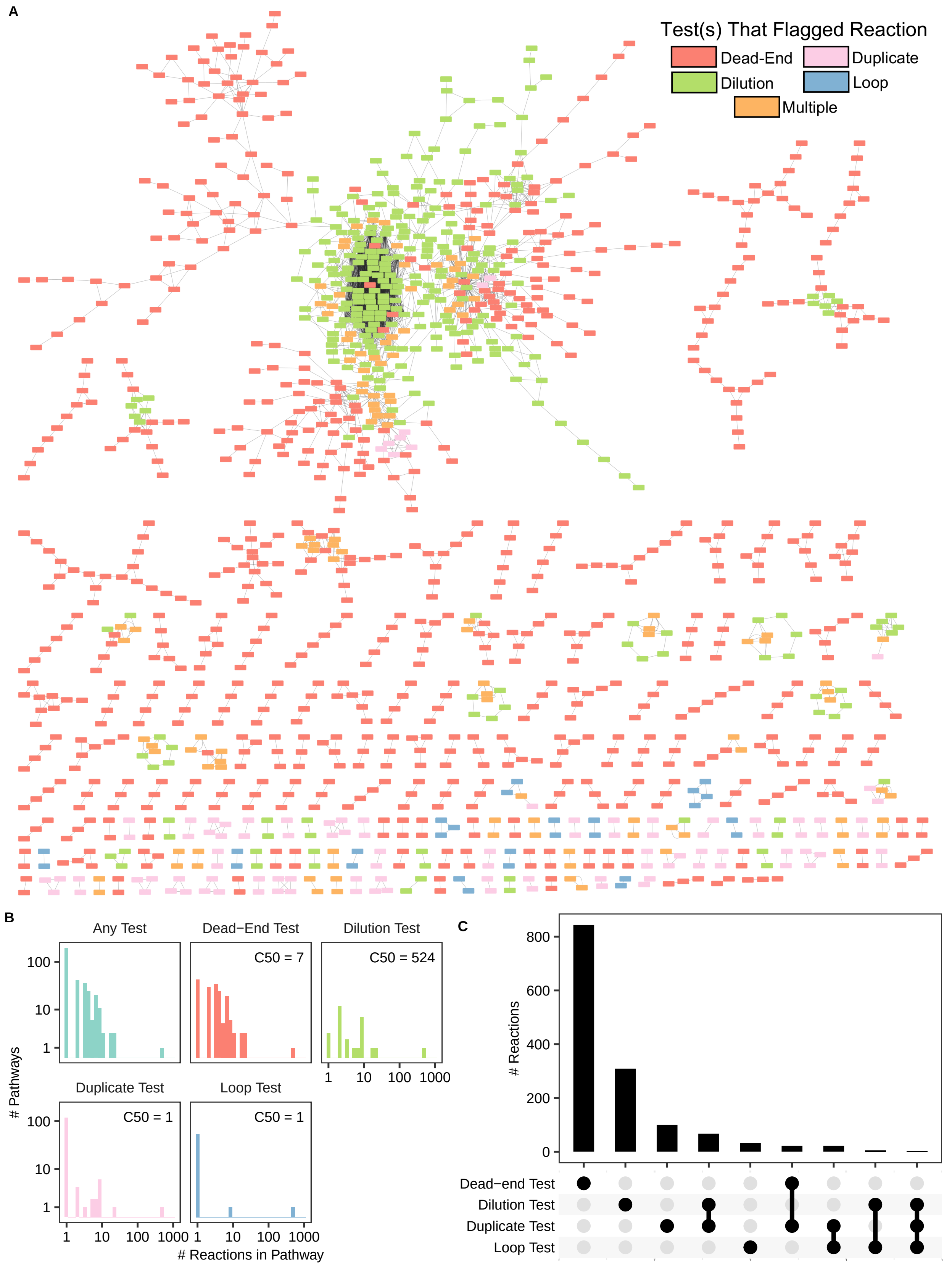


**Supplementary Figure 3. Overview of reactions in iML1515 flagged by one or more tests in**

**MACAW.** (A) Each node represents a single reaction; see Methods for explanation of how reactions were connected. The color of each node indicates which test(s) the reaction was flagged by. (B) Distributions of numbers of reactions in each connected component ("pathway") shown in (A) for all pathways or only pathways containing at least one reaction flagged by the specified test. The C50 of each distribution is the median number of reactions flagged by each test weighted by the total number of reactions in the same pathway as each reaction. (C) UpSET plot showing number of reactions flagged by each observed combination of tests in MACAW.

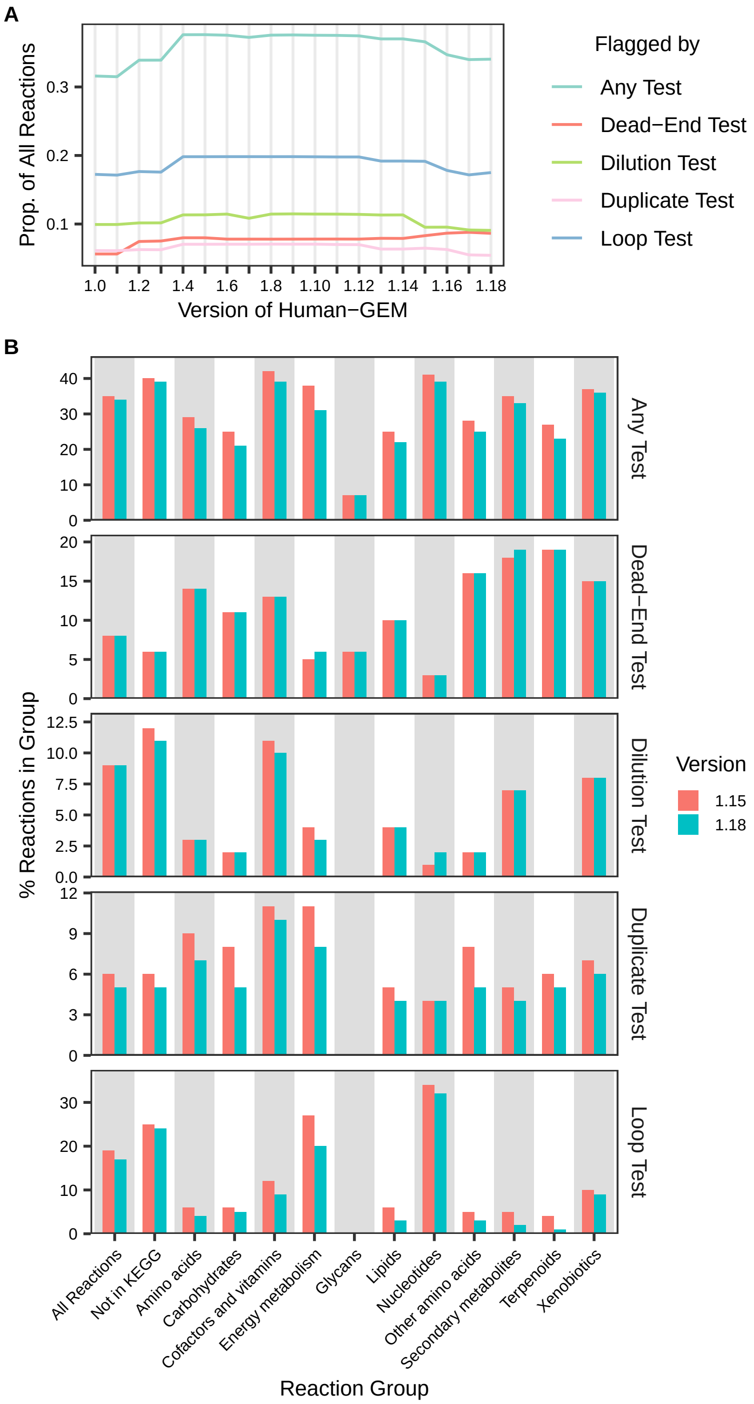


**Supplementary Figure 4. Comparison of test results in different versions of Human-GEM.** (A) Proportions of reactions flagged by tests across all versions of Human-GEM. (B) Proportions of reactions flagged by tests in versions 1.15 and 1.18 of Human-GEM. Reactions grouped according to the second-highest levels of the KEGG functional ortholog hierarchy containing the genes associated with each reaction; see "Methods". Individual reactions may be associated with more than one group of KEGG functional orthologs. Names of KEGG functional ortholog groups have been abbreviated.

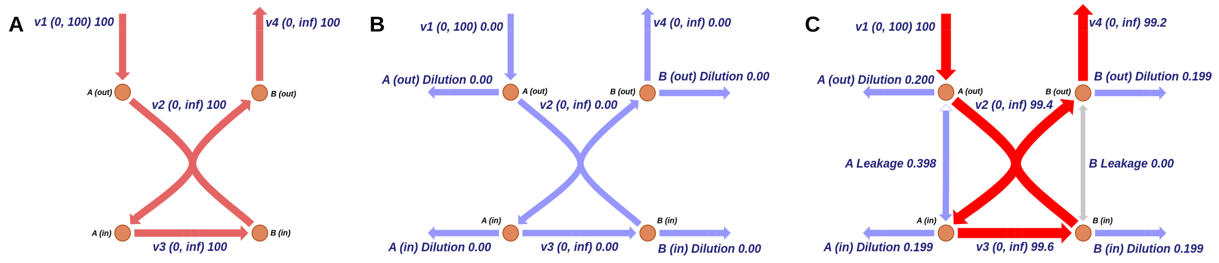


**Supplementary Figure 5. "Leakage" reactions prevent the dilution test from flagging unproblematic antiport reactions.** (A) Toy model with two metabolites that can exist in two different compartments: "out" and "in", and move between them via the antiport reaction v2. Numbers in parentheses next to reaction labels are the minimum and maximum allowed fluxes through that reaction. Numbers following reaction bounds are the optimal fluxes through each reaction when maximizing flux through v4. (B) Same as (A) except one dilution reaction has been added for each metabolite whose flux is constrained to be exactly equal to 0.1% of the sum of the absolute values of all other fluxes that involve that metabolite. (C) Same as in (B) except one leakage reaction has been added connecting each pair of metabolites that exist in two separate compartments. Fluxes through leakage reactions are constrained to be between -1 and 1.
