## Supplementary material for "Semi-Automatic Detection of Errors in Genome-Scale Metabolic Models": Table 1

**Tables**

|  | **MACAW** | **MEMOTE** | **ErrorTracer** |
| --- | --- | --- | --- |
| **Duplicate Reactions** |  |  |  |
| Completely identical | all | some | all |
| Same metabolites, different coefficients | all | some | some |
| Same metabolites, different directions | all | some | some |
| Same metabolites, different reversibility | all | some | some |
| Same genes, everything else potentially different | some | all | some |
| Same metabolites except for redox metabolites | all | none | none |
| **Dead-End Metabolites and Reactions** |  |  |  |
| Metabolites that can only be produced | all | some | all |
| Metabolites that can only be consumed | all | some | all |
| Reactions that involve those metabolites | all | none | all |
| Reactions upstream or downstream of those | all | none | some |
| Accounts for effectively irreversible reactions? | yes | no | yes |
| **Thermodynamically Infeasible Cycles** |  |  |  |
| Blocked by dilution constraints | all | some | some |
| Internal loops (type III extreme pathways) | all | all | all |
| Energy-generating cycles | some | all | some |

**Table 1. Comparison of MACAW to similar previously published tools.** Effectively irreversible reactions are reversible reactions that have at least one product or reactant that can only be consumed by every other reaction that it participates in or only be produced by every other reaction that it participates in. See “Methods” for definition of dilution constraints. Internal loops (type III extreme pathways) are groups of reactions that can sustain non-zero steady-state fluxes while no exchange reactions (reactions that represent consumption of nutrients from or secretion of metabolites into the environment of a cell) have non-zero steady-state fluxes. Energy-generating cycles are sets of reactions that can sustain non-zero steady-state fluxes while the only exchange reactions with non-zero steady-state fluxes are those for energy-currency metabolites such as ATP and NAD(H).
